# Cancer SIGVAR: A semi-automated interpretation tool for germline variants of hereditary cancer-related genes

**DOI:** 10.1101/2020.04.15.042283

**Authors:** Hong Li, Shuixia Liu, Shuangying Wang, Quanlei Zeng, Yulan Chen, Ting Fang, Yi Zhang, Ying Zhou, Yu Zhang, Kaiyue Wang, Zhangwei Yan, Cuicui Qiang, Meng Xu, Xianghua Chai, Yuying Yuan, Ming Huang, Hongyun Zhang, Yun Xiong

## Abstract

The American College of Medical Genetics and Genomics and the Association for Molecular Pathology published guidelines in 2015 for the clinical interpretation of Mendelian disorder sequence variants based on 28 criteria. ClinGen Sequence Variant Interpretation (SVI) Working Groups have developed many adaptations or refinements of these guidelines to improve the consistency of interpretation. We combined the most recent adaptations to expand the criteria from 28 to 48 and developed a tool called Cancer SIGVAR to help healthcare workers and genetic counselors interpret the clinical significance of cancer germline variants, which is critical for the clinical diagnosis and treatment of hereditary cancer. Our tool can accept VCF files as input and realize fully automated interpretation based on 21 criteria and semi-automated interpretation based on 48 criteria. We validated our tool on the ClinVar and CLINVITAE benchmark databases for the accuracy of fully automated interpretation, achieving an average consistency for pathogenic and benign assessment up to 93.40% and 82.54%, respectively. We compared Cancer SIGVAR with a similar tool, InterVar, and analyzed the main differences in criteria and implementation. In addition, to verify the performance of semi-automated interpretation based on 48 criteria, we selected 911 variants from two benchmark databases and reached an average classification consistency of 98.35%. Our findings highlight the need to optimize automated interpretation tools based on constantly updated guidelines.

## Introduction

With the rapid development of next-generation sequencing (NGS) technologies, the use of genetic test results to guide clinical diagnosis and medication has received increasing attention (Biesecker & Green, 2014). However, with hundreds to thousands of variants per individual across genome sequencing, accurately interpreting the clinical significance of variants and their relationship with clinical symptoms is a key component of accurate diagnosis that has a serious impact on clinical care (Luo et al., 2019). To standardize the interpretation of variants, the American College of Medical Genetics and Genomics and the Association of Molecular Pathologists (ACMG-AMP) published a joint consensus recommendation that provided a set of criteria and specific terms to interpret and describe the clinical significance of variants in Mendelian disease (Richards et al., 2015).

The ACMG-AMP guidelines provide a theoretical basis and action directions for variant interpretation that can to some extent improve the consistency and reliability of the interpretation of genetic variations, but there are still certain limitations. First, interpreters have different abilities to obtain literature and database information, extract phenotypic data and understand the classification criteria. Additionally, they use different databases and prediction algorithms (Amendola et al., 2016). Therefore, there are inconsistencies in the classification of the same variant by different laboratories, even among professional genetic testing laboratories. For example, surveys have shown that the application of the ACMG guidelines by nine US laboratories classifies 99 variants, with only 34% consistency in the initial classification, of which 22% of the differences may affect clinical decision-making (e.g., LP/P vs. VUS/LB/B) (Amendola et al., 2016). Second, although the main goal of the ACMG-AMP guidelines is to standardize the interpretation of variation, there is no clear description of the use of each criterion, and some lack specificity, which leads to interpretation differences (Kelly et al., 2018; Nykamp et al., 2017). To further promote the consistency of interpretation, the Clinical Genomic Resource (ClinGen) Sequence Variant Interpretation (SVI) Working Group has provided more detailed guidance and recommendations on the application of some criteria in the ACMG-AMP guidelines, including the use of PVS1 (Abou Tayoun et al., 2018), the level of evidence strength of some criteria (e.g., PP1, PS2/PM6, PS4, PM4, etc.) (Luo et al., 2019), setting the allele frequency (AF) threshold (BA1, BS1, PM2) (Gelb et al., 2018; Oza et al., 2018), etc. In addition, the SVI Working Groups have successively published optimized guidelines for specific diseases (such as hearing loss (Oza et al., 2018), RASopathy phenotype (Gelb et al., 2018), etc.) or specific genes (such as *PTEN* (Mester et al., 2018), *CDH1* (Lee et al., 2018), *PAH* (Zastrow et al., 2018), *MYH7* (Kelly et al., 2018), etc.). Adaptation and optimization of the guidelines have increased the consistency of variant interpretation.

With the continuous refinement of the ACMG-AMP guidelines, the consistency of interpretation has increased, as has the complexity of the evaluation criteria. An inconsistent understanding of variants, such as co-segregation and *de novo* status, and the complexity and error-proneness of database retrieval are important challenges for the accurate interpretation of variants. To simplify the manual interpretation process, the variant interpretation tool generated by a combination of guidelines and automated procedures has gradually become an industry trend. Some easy-to-use automated variant interpretation tools based on the 2015 ACMG-AMP guidelines, such as InterVar (Li & Wang, 2017), the ClinGen Pathogenicity Calculator (Patel et al., 2017) and CardioClassifier (Whiffin et al., 2018), have been developed to generate repeatable criteria for each variant and to help to interpret and understand the clinical significance of germline variants. The use of automated tools solves the complicated nature of the manual interpretation of database retrieval and to some extent improves the repeatability of interpretation. However, automated interpretation tools should be optimized with continuously adapted guidelines. The CardioVAI automated interpretation tool combines CMP-EP’s adaptation with the 2015 ACMG-AMP guideline and reaches a high consistency of 83% pathogenic and 97.08% benign for CVD-related gene variants (Nicora et al., 2018). Studies have shown that adding reasonable adaptation and refinement of the ACMG-AMP guidelines to the automated interpretation tool can increase the accuracy of automated variant classification and simultaneously improve the efficiency of variant interpretation.

Since the ACMG-AMP guidelines were published, they have been widely adopted and adapted to many diseases, such as hereditary cardiomyopathy (Kelly et al., 2018), cardiovascular disease (Nicora et al., 2018) and hereditary breast cancer (Maxwell et al., 2016). Cancer is one of the most important health issues globally and a possible application area recommended by ACMG-AMP. In 2018, estimations indicate approximately 18.1 million new cases of cancer and 9.6 million cancer deaths worldwide (Bray et al., 2018). The occurrence and development of cancer are influenced by both genetic and environmental factors, with approximately 5-10% of cancers being caused by genetic factors (Nagy, Sweet, & Eng, 2004). Mutations in cancer-susceptibility genes can, to a certain extent, increase the risk of cancer in individuals. Since 2005, the application of gene detection technology in the field of cancer diagnosis has gradually reversed the survival situation of cancer patients. For example, more than 5 million people in the United States have undergone genetic testing, which has significantly improved the cure rate of some major chronic diseases; for example, the incidence of familial bowel cancer has been reduced by 90%, and the mortality rate has been reduced by 70% (Rebecca Siegel, 2019). The essential premise of genetic testing in guiding accurate diagnosis and treatment is the accurate interpretation of variants, so it is very meaningful to develop automated interpretation tools for hereditary cancer genes. There is currently no automated interpretation tool containing the latest adaptations of the ACMG-AMP guidelines and specific to cancer susceptible genes. In terms of classification logic, the current automated interpretation tools are not sufficiently perfected.

To automate, standardize and professionalize the interpretation of hereditary cancer-associated germline variants, we have developed Cancer SIGVAR. This automated tool is based on the adaptation of ACMG-AMP guidelines and the reasonable handling of the logical relationship between criteria. It is appropriate for the interpretation of 97 susceptible genes in hereditary cancer. This tool not only improves the accuracy of variant interpretation but also realizes automated interpretation, which allows it to produce fast, repeatable and reliable results for the interpretation of genetic variants.

## Material and Methods

### Variant annotation generation

We developed annotation software to annotate genetic variants using various biological databases and the standard variant type and name based on SO (Eilbeck et al., 2005) and HGVS (den Dunnen et al., 2016), respectively. Users can configure the annotation resources in VCF format. In this study, we annotated variants based on Genome Reference Consortium Human Genome Build 37 (GRCh37), and the annotation resources configured for automated interpretation included (1) populational data, such as NHLBI Exome Sequencing Project (ESP6500), Exome Aggregation Consortium (ExAC), Genome Aggregation Database (gnomAD), and in-house databases; (2) predicted data, such as dbNSFP (v3.5a) (Liu, Jian, & Boerwinkle, 2011; Liu, Wu, Li, & Boerwinkle, 2016), dbscSNV (1.1) (Jian, Boerwinkle, & Liu, 2014), DANN (Quang, Chen, & Xie, 2015), GERP (Davydov et al., 2010); and (3) databases: ClinVar database and in-house database.

### Criteria assignment and scoring system

Cancer SIGVAR expanded 28 criteria to 48 criteria by combining the most recent guidelines, which were adapted based on considerations of gene specificity, disease specificity and level of evidence strength (Abou Tayoun et al., 2018; Gelb et al., 2018; Karczewski et al., 2019; Lee et al., 2018; Luo et al., 2019; Mester et al., 2018; Oza et al., 2018). A detailed description of these 48 criteria is shown in Figure 1. Among them, 21 criteria received automated scores, 17 criteria received semi-automated scores, and 10 criteria were manually scored. The implementation for the 48 criteria in Cancer SIGVAR are shown in Supporting Information Table S1.

**Figure 1.**
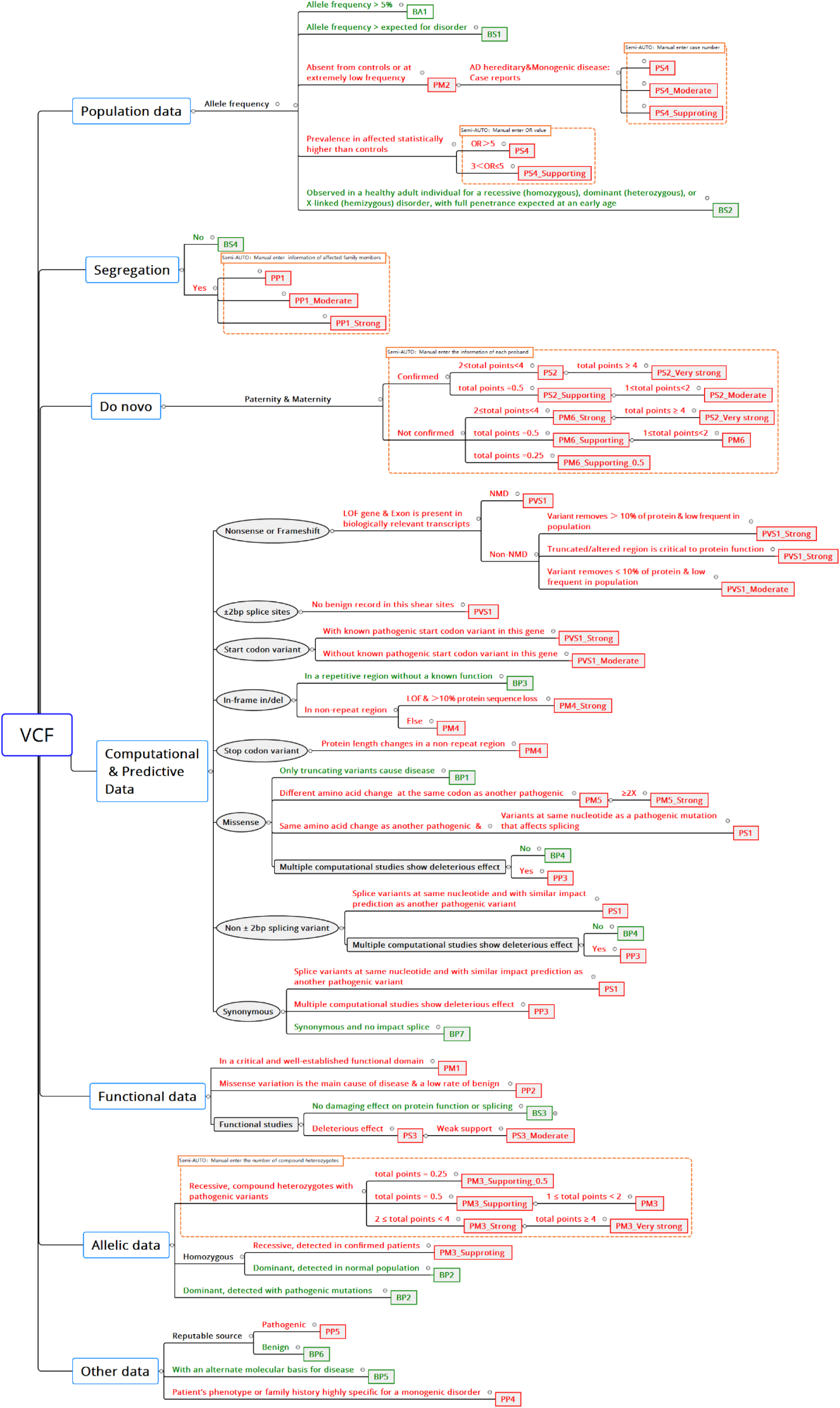
Illustration of the 48 Criteria used in Cancer SIGVAR

To realize automatic assignment and standardization of the above criteria, Cancer SIGVAR integrates the ACMG-AMP proposed public knowledge base and further considers key specifications, such as phenotypic specificity (inheritance, penetrance, disease mechanism), AF thresholds, incidence rate and validity of functional assays.

For PVS1, we confirmed the mechanism of LOF (loss of function) and nonsense-mediated mRNA decay (NMD) in each gene according to phenotypic specificity and Mutation Taster prediction. The gene functional domain and protein repeat region were obtained from InterPro and RefSeq & UniProt, respectively, which are applicable to PVS1, PM1, PM4 and BP3. In silico tools such as dbscSNV, DANN, GREP++, Polyphen2, SIFT and Mutation Taster are recommended for implementing PP3, BP4 and BP7. Polyphen2, SIFT and Mutation Taster are used for protein function prediction. dbscSNV_RF and dbscSNV_ADA scores were used to splice impact predictions, while GREP++ and DANN are useful for the prediction of evolutionary conservation and deleteriousness, respectively. See Supporting Information Table S1 for the detailed usage of prediction software for different criteria. As a reference for the bona fide pathogenic and benign variants necessary for PS1, PM5, PP5 and BP6, we used the ClinVar repository (version 2019-04-03) and the in-house database. We filtered ClinVar with sole interpretation and review status ≥ 2 stars to ensure the accuracy and reliability of variant interpretation and combined it with the in-house database by deleting the same variants to form an integrated database.

Our implementation of population frequency criteria, i.e., PM2, BA1, and BS1, relies on five different databases: gnomAD, 1000 Genomes Project, ExAC, ESP6500 and our in-house database. In addition, BS2 and BP2 are automated by using individual population research from gnomAD. Corresponding criteria are assigned based on the AF of variants in the database; specific thresholds and rules are shown in Supporting Information Table S1.

Failure to achieve full-automated scoring for all criteria is due to a lack of literature or patient-specific information (such as co-segregation of genotype-phenotype, effective functional experiments, the number of probands and the *de novo* status of variants). Notably, we combine built-in calculation programs with manually entered parameters to achieve semi-automated scoring of PS2, PM6, PM3 and PS4 for 17 criteria, which can reduce manual workload and possible errors in determining the level of evidence. For example, we have set up a scoring system for PM3 based on optimization of ClinGen Expert and the ACMG-AMP guidelines. After the information of each proband is manually entered, Cancer SIGVAR will automatically identify the corresponding point value of each proband according to the rules in Supporting Information Table S2 and calculate the total points to determine the final level of the criteria. For the other 10 criteria requiring manual assignment, we also formulated the corresponding interpretation standards. For details, see Supporting Information Table S1.

Finally, we optimize the conflict handling rules. First, logical complementarity between pathogenic and benign evidence has been set to reduce the interpretation of contradictions. Second, when different criteria for functional evidence coexist, we set a rule that PP3 and BP4 cannot be applied in combination with PVS1 to avoid the double use of evidence. Finally, the tolerance value is set when pathogenic evidence and benign evidence coexist. The detailed decision logic is shown in Supporting Information Table S3.

### Benchmark datasets

We selected four benchmark datasets to evaluate the accuracy of the variant interpretation of Cancer SIGVAR. CLINVITAE is a clinical variation database that is collected and integrated from public resources. The data sources include the ClinVar database and other open laboratory databases. This database has been used as a benchmark dataset multiple times (Li & Wang, 2017; Nicora et al., 2018). We selected 45,463 variants of sole interpretation in 97 cancer-susceptibility genes from the CLINVITAE version 2019-09-04 database (https://clinvitae.invitae.com/) as Dataset 1. ClinVar is a public database that is freely accessible and does not modify or evaluate the accuracy of submitted interpretations (Harrison et al., 2016). Increasing research indicates that data in ClinVar may also have errors (Shi et al., 2016). ClinVar assigns review status (0 to 4 stars) to variants based on all submissions; the more stars there are in the variant review status, the more reliable the variant interpretation is (Harrison et al., 2016). Considering the accuracy and reliability of the interpretation results of variants, we screened 26,065 variants of sole interpretation and review status ≥ 2 stars in 97 cancer-susceptibility genes from the ClinVar database version 2019-04-03 (https://www.ncbi.nlm.nih.gov/clinvar/) as Dataset 2. The accuracy of the automated interpretation of Cancer SIGVAR was evaluated by comparing the results of the pathogenicity classification with the results of two benchmark databases. Database 3 is a total of 731 unique variants collected by the National Institutes for Food and Drug Control and represents a standard database for genetic variant interpretation in China. We selected 180 variants from the BGI in-house database that cover most of the evidence as Dataset 4. These two in-house datasets were used as benchmarks to evaluate the accuracy of the semi-automated interpretation based on all 48 criteria.

## Results

### Summary of the interpretation procedure

Cancer SIGVAR is a semi-automated interpretation tool for 97 susceptible genes (Supporting Information Table S4) in hereditary cancer, and the workflow is shown in Figure 2. Cancer SIGVAR mainly consists of four major steps: (1) Integrated annotation: we use self-developed annotation software to extract and annotate the relevant parameters of the variants and then use the population database, automated prediction software and other in-house databases to supplement and annotate the other parameters. (2) Automated scoring: Cancer SIGVAR offers a fully automated score for 21 criteria based on annotated information at the first step and then output all relevant criteria and a preliminary variant interpretation. (3) Manual adjustment: Semi-automated scoring of 17 criteria involving PS2, PM6, PM3 and PS4 combines built-in calculation programs with manually entered parameters. The manual assignment of 10 criteria is based on user domain knowledge. (4) The fourth step is automated classification, realizing fully automated interpretation based on 21 automated scoring criteria and semi-automated interpretation based on all 48 criteria including manual adjustment. Then, Cancer SIGVAR outputs the final interpretation results.

**Figure 2.**
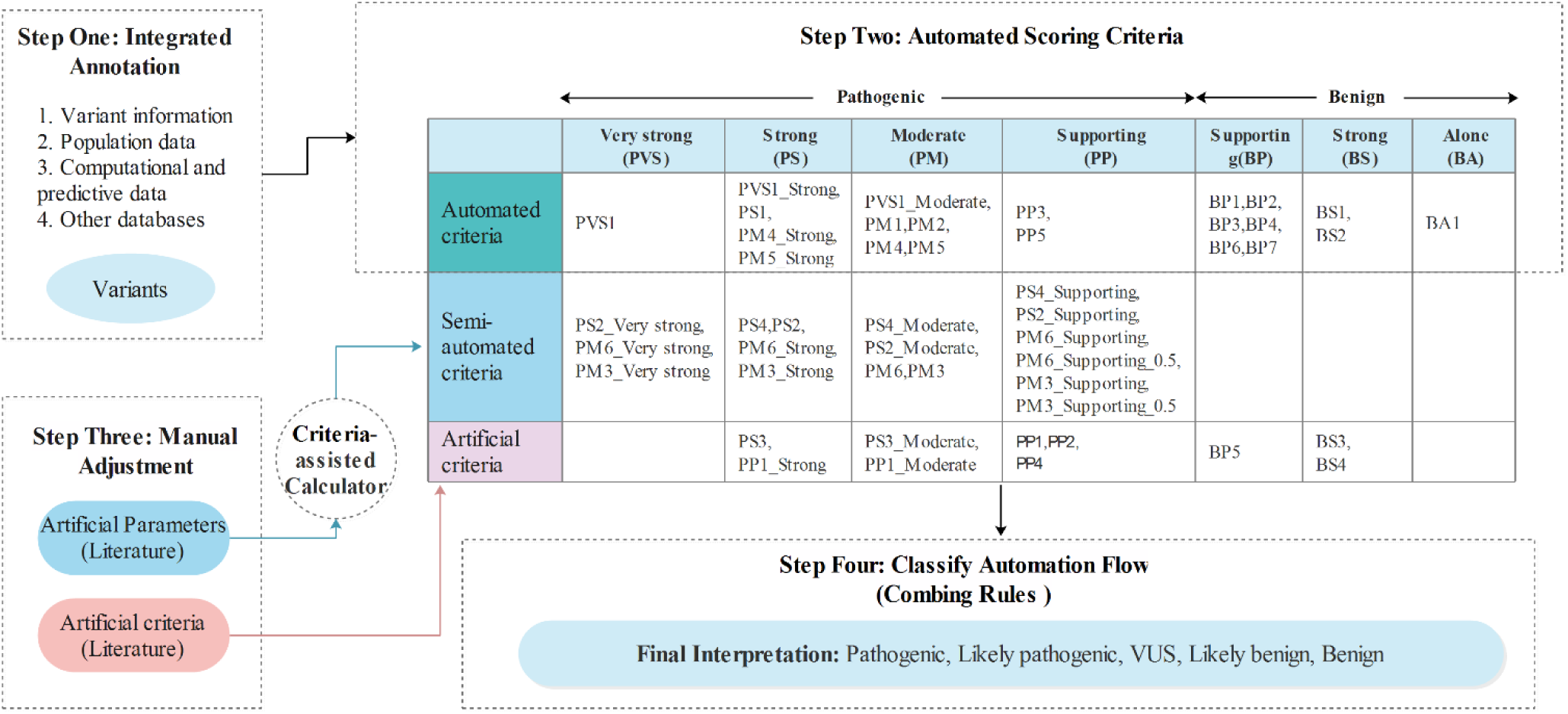
The Workflow of Cancer SIGVAR

As an example, we illustrate the procedure for using Cancer SIGVAR by interpreting the missense variant *BRCA1* c.181T>G (p. Cys61Gly) (GRCh37 coordinate) in exon 4, which associates with hereditary breast cancer and ovarian cancer. In the first step, the basic parameters of the variant are automatically extracted and annotated by self-developed annotation software. Then, Cancer SIGVAR outputs the automated criteria based on the integrated annotation. The AFs of this variant in gnomAD and ExAC are less than 0.00008 and without any AF ≥ 0.005 in all population databases, so PM2 is assigned. In addition, multiple computer simulation calculations predict the variant is harmful, and two different pathogenic amino acid changes of p. Cys61Tyr are determined, and thus PP3 and PM5_Strong are applied. For this variant, the conclusion of the ClinVar dataset is pathogenic, so PP5 is assigned. According to the rules, the variant falls into the class of ‘‘Likely pathogenic’’. For a web presentation of the automated interpretation, see supporting information Figure S1A. In the third step, we make manual adjustments through literature retrieval. We first enter the relevant parameters manually with an OR value of 26 and 95% CI of 6.1 to 113.7, which were confirmed through the literature and case-control studies (Brozek et al., 2011). Then, built-in calculation tools assist with semi-automated scoring and output criteria PS4. In addition, experimental studies have shown that this missense variant disrupts proper function of the *BRCA1* protein (Bouwman et al., 2013; Brzovic, Meza, King, & Klevit, 1998; Drost et al., 2011; Towler et al., 2013), so PS3 is artificially assigned. The variant is co-segregated with disease (Friedman et al., 1994), so PP1 is manually assigned. The corresponding web interface for manual adjustment is shown in Supporting Information Figure S1B. In the fourth step of Cancer SIGVAR, the classification automation process will integrate all criteria (PM2, PM5_Strong, PP3, PP5, PP1, PS3, PS4) and reinterpret the variant as “Pathogenic” on the basis of “≥2 Strong (PS1–PS4)”; the web interface is shown in Supporting Information Figure S1C. This procedure illustrates how to use the automated interpretation and manual adjustment of Cancer SIGVAR to derive a final interpretation for genetic variants.

### Automated interpretation of cancer-susceptibility variants is highly consistent with ClinVar

To assess the performance of Cancer SIGVAR, we selected 26,065 sole interpreted variants in 97 cancer-susceptibility genes from the filtered ClinVar database. The types of these variants and the corresponding number of each type are shown in Supporting Information Figure S2A. Among them, 7440 were interpreted as pathogenic (P) or likely pathogenic (LP), 11,497 were interpreted as a variant of unknown significance (VUS), and 7128 were interpreted as benign (B) or likely benign (LB). We then reinterpreted these variants using the automated interpretation program in Cancer SIGVAR (Table 1). The results show that the consistency of the pathogenic (P/LP) and benign (B/LB) categories between Cancer SIGVAR and ClinVar is 94.74% (7049/7440) and 83.46% (5949/7128), respectively, indicating that the automated interpretation of Cancer SIGVAR is highly concordant with the results of ClinVar and reaches high accuracy and specificity. It should be noted that these explanations made by Cancer SIGVAR are based on the 21 criteria for automated scoring in the second step, without the manual assignment in the third step. If manual assignment is added, it becomes more accurate.

**Table 1.** Illustration of Automated Interpretation of Pathogenic and Benign Variants Annotated in ClinVar.

| Cancer SIGVAR<br>(Automated Interpretation) | ClinVar |  |  |
| --- | --- | --- | --- |
|  | Pathogenic or<br>Likely Pathogenic | VUS | Benign or Likely<br>Benign |
| Pathogenic | 6305 | 0 | 0 |
| Likely pathogenic | 744 | 45 | 0 |
| VUS | 390 | 11378 | 1179 |
| Likely benign | 0 | 67 | 5336 |
| Benign | 1 | 7 | 613 |
| Sum of five tiers | 7440 | 11497 | 7128 |
| Pathogenic or likely<br>pathogenic | 7049 (94.74%) | 45 (0.39%) | 0 (0%) |
| Benign or likely benign | 1 (0.013%) | 74 (0.64%) | 5949 (83.46%) |

For the differences in interpretation between ClinVar and Cancer SIGVAR, we conducted a more detailed analysis of 1 (0.013%) variant classified as pathogenic by ClinVar and benign by Cancer SIGVAR. It is a frameshift variant NM_000057.4: c.2923del in the *BLM* gene. In ClinVar, the review status level of this variant is two stars, and it was interpreted as pathogenic by five different agencies. In the ESP6500 database, the AF of this variant is 0.1504; therefore, Cancer SIGVAR’s automated scoring assigned BA1, of which the final interpretation is benign. It should be noted that the frequency of this variant in the ESP6500 database is incorrect. Considering that this is an objective problem within public databases, the interpretation requires manual adjustment.

### Automated interpretation of cancer-susceptibility variants is highly consistent with CLINVITAE

To further analyze the accuracy of the automated interpretation of Cancer SIGVAR, we selected 45,463 cancer-specific genetic variants (types of variants are shown in Supporting Information Figure S2B) from the CLINVITAE database that belong to the 97 cancer-susceptibility genes. Unlike ClinVar, which did not modify the submitted interpretations, CLINVITAE integrates the collected variant interpretation data, and they are interpreted by the INVITAE team. Among these 45,463 variants, 5775 and 9404 variants were classified as pathogenic or likely pathogenic and benign or likely benign, respectively (Table 2). Among them, Cancer SIGVAR also classified 5316 (92.05%) and 7676 (81.62%) as P/LP and B/LB, respectively. This analysis once again proves the high consistency between the automated interpretations of Cancer SIGVAR and the classifications made by specialists. The variant NM_000057.4 (*BLM*): c.2923del, which was described above, was classified as pathogenic by CLINVITAE and classified as benign by Cancer SIGVAR due to data error in the ESP6500 database.

**Table 2.**
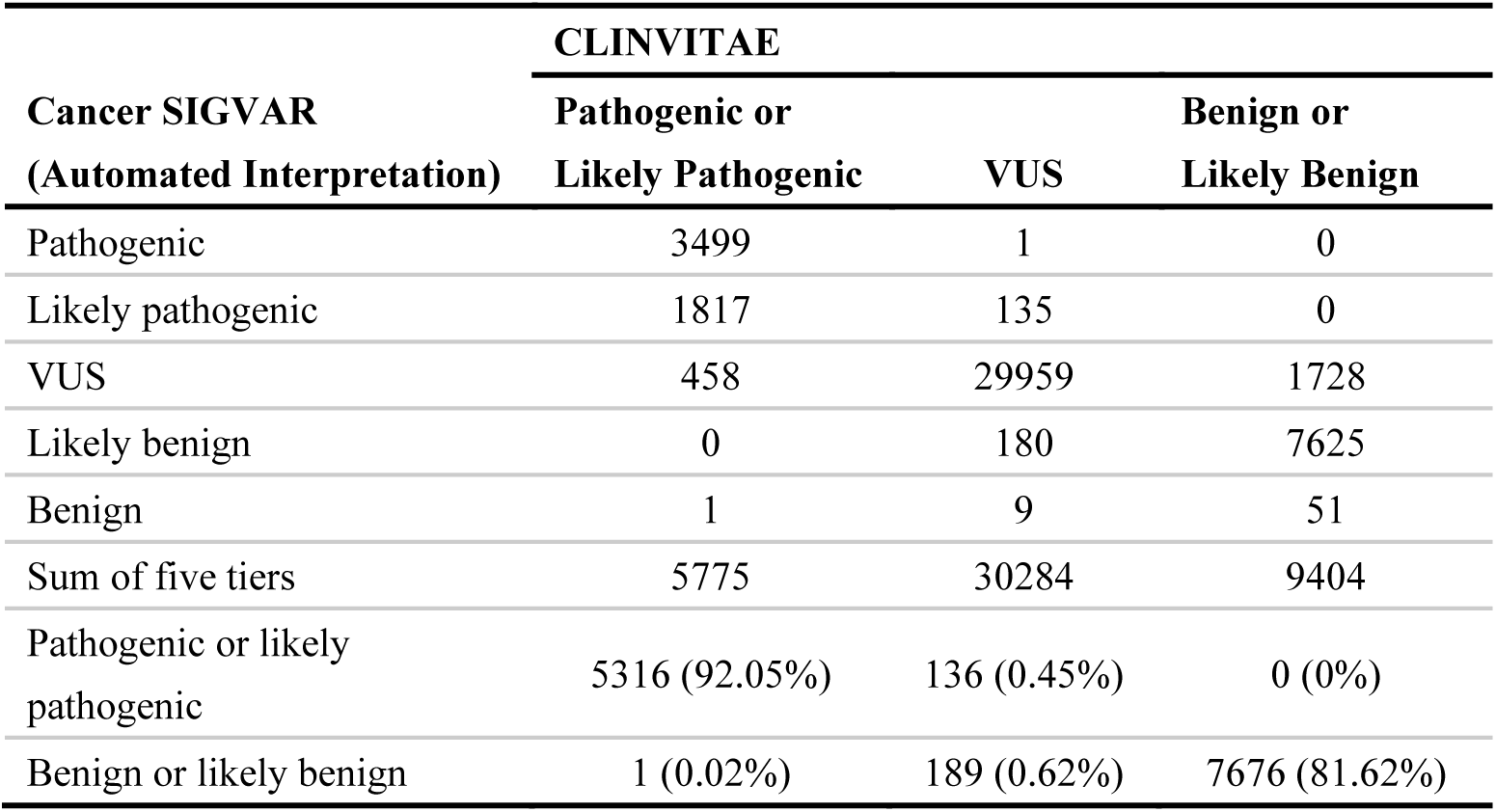
Illustration of Automated Interpretation of Pathogenic and Benign Variants Annotated in CLINVITAE.

### Comparative analysis with InterVar

We downloaded a similar ACMG-based automated interpretation tool InterVar (version py 2.0.2 20190327) and analyzed the above ClinVar and CLINVITAE benchmark databases using the default settings. We then conducted a comparative analysis of the differences between Cancer SIGVAR and InterVar in automated interpretation. The consistency results with ClinVar are shown in Table 3. For the P/LP category, the consistency rate of Cancer SIGVAR is 94.74% (7049/7440), which is higher than that of InterVar, 92.22% (6861/7440). However, for the B/LB categories, the consistency rate of Cancer SIGVAR is 83.46% (5949/7128), which is lower than the 88.73% (6325/7128) of InterVar. For all variant classifications, the overall consistency of Cancer SIGVAR is 93.52% (24376/26065), which is higher than that of InterVar, 87.03% (22685/26065). The consistency results with CLINVITAE are shown in Supporting Information Table S5. The overall consistency of the automated interpretation of Cancer SIGVAR is 94.47% (42951/45463), which is higher than that of InterVar 83.70% (38052/45463).

**Table 3.** Comparison of Variant Automated Interpretation consistent with ClinVar by InterVar and Cancer SIGVAR.

| Clinical Significance | ClinVar | InterVar (Automated Interpretation) | Cancer SIGVAR (Automated Interpretation) |
| --- | --- | --- | --- |
| Pathogenic or likely pathogenic | 7440 | 6861 (92.22%) | 7049 (94.74%) |
| Benign or likely benign | 7128 | 6325 (88.73%) | 5949 (83.46%) |
| VUS | 11497 | 9499 (82.62%) | 11378 (98.96%) |
| Sum of five tiers | 26065 | 22685 (87.03%) | 24376 (93.52%) |

To determine the reasons for the differences in the automated interpretation between Cancer SIGVAR and InterVar, we further analyzed the difference in criteria assignment between these two tools for pathogenic and benign variants from the ClinVar database. As shown in Figure 3A, for the P category in ClinVar, the consistency in the interpretation of Cancer SIGVAR is higher than that of InterVar (91.18% vs 90.54%). The relative consistency (LP/P and P/P) of classification in Cancer SIGVAR is higher (96.87% vs 93.06%). As shown in Figure 4B, the main reason is the difference in the use of PVS1 and its downgraded criteria PVS1_Strong. The total criteria frequency difference in P/LP (Figure 3C) also illustrates this. According to the 2015 ACMG-AMP guidelines and the adaptation of PVS1 by ClinGen (Abou Tayoun et al., 2018), PVS1 criteria are mainly applied to initiation codon, spliceDonor, spliceAcceptor, nonsense and frameshift types of the SNV or INDEL variant (CNV is currently not implemented in Cancer SIGVAR), and whether nonsense and frameshift types are predicted to undergo NMD will lead different levels of PVS1 criteria, so we divide the variants in ClinVar into four categories as shown in Figure 4A. The results show that the consistency in the automated interpretation of Cancer SIGVAR with ClinVar is significantly higher than that of InterVar. In particular, in the case of non-NMD, compared to InterVar, the consistency of pathogenic interpretation was improved by 54.42% (from 115 to 306). Figure 4B shows the differences in criteria activation between Cancer SIGVAR and InterVar for those variants that were correctly interpreted by Cancer SIGVAR but not by InterVar. The results show that Cancer SIGVAR sets up a hierarchy of criteria and uses a more comprehensive PVS1 classification, which helps to improve the accuracy of pathogenic interpretation.

**Figure 3.**
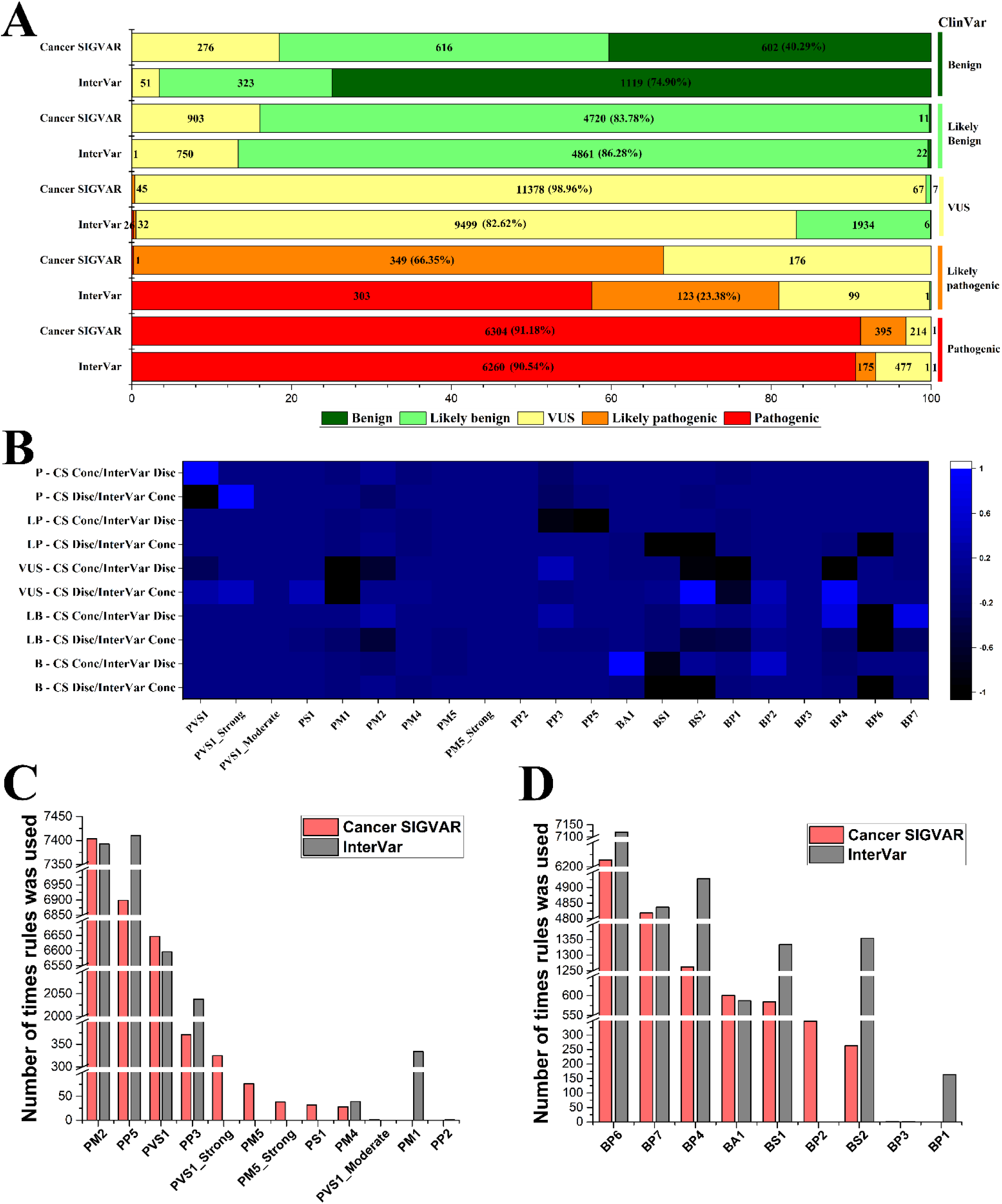
(A). The automated interpretations of Cancer SIGVAR and InterVar in different categories of ClinVar. (B). Differential criteria heatmap between Cancer SIGVAR and InterVar on different categories of variants.CS Cons/InterVar Disc means those variants that were correctly interpreted by Cancer SIGVAR but not by InterVar. CS Disc/InterVar Cons means those variants that were correctly interpreted by InterVar but not by Cancer SIGVAR. Higher values indicate that a criterion has been triggered more by Cancer SIGVAR, while lower values show criteria triggered more times by InterVar. (C). Frequency of pathogenic criteria applied in interpretation of Cancer SIGVAR and InterVar for 7440 variants of pathogenic or likely pathogenicity categories of ClinVar. (D). Frequency of benign criteria applied in interpretation of Cancer SIGVAR and InterVar for 7128 variants of benign or likely benign categories of ClinVar

**Figure 4.**
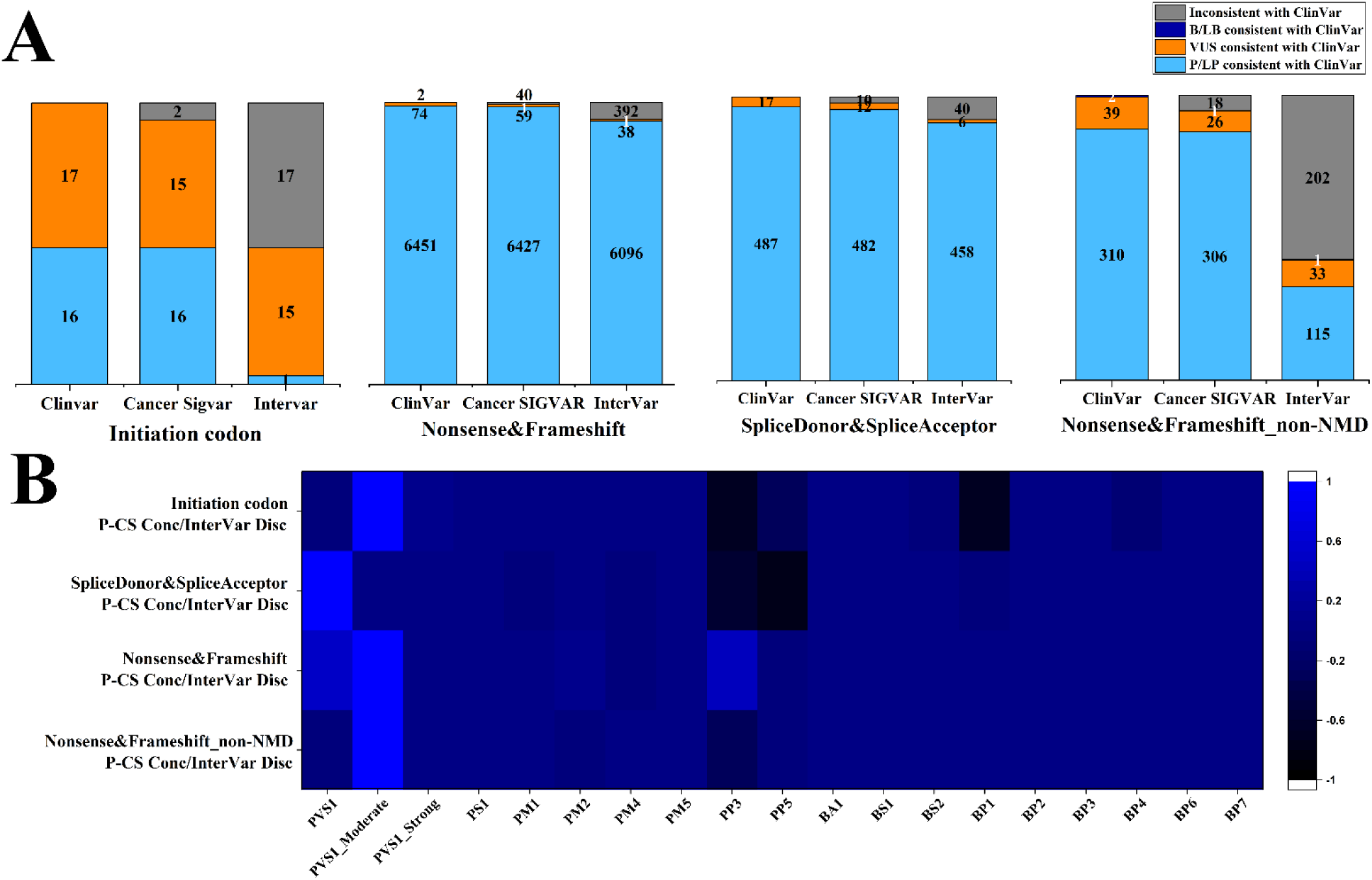
(A). For different types of variants, compare the consistency of Cancer SIGVAR and InterVar automated interpretation results with ClinVar. (B). Differential criteria heatmap between Cancer SIGVAR and InterVar on different types of pathogenic variants: each criterion is associated to the difference of means in the number of times it is activated by Cancer SIGVAR and by InterVar for those variants that were correctly interpreted by Cancer SIGVAR but not by InterVar. Higher values indicate that a criterion has been triggered more by Cancer SIGVAR, while lower values show criteria triggered more times by InterVar.

For the likely pathogenic variants in ClinVar, the consistency of Cancer SIGVAR is higher than that of InterVar (66.35% vs 23.38%), but the relative consistency of LP/P is lower than that of InterVar. InterVar automatically interprets many LP variants as P. As shown in Figure 3B-C, the main difference in the use of pathogenic criteria is PP3 and PP5. To ensure the comprehensiveness and accuracy of prediction as well as to avoid overprediction, more prediction software with different strategies is used for different types of variation, different criteria of the same type were avoided, and we also strictly distinguish between null and 0 for the score of the prediction algorithms. Therefore, the frequency of use of the PP3 and BP4 criteria is significantly lower than that of InterVar. Combining the opinions of the ClinGen SVI team and the ACMG working group on the use of PP5 and BP6 (Biesecker & Harrison, 2018; Richards et al., 2015). We apply the PP5 and BP6 criteria in Cancer SIGVAR under stricter conditions to ensure the reliability of the criteria. Details can be found in Supporting Information Table S1. The strict use of PP5 and BP6 reduces errors in variant classification. As shown in Figure 3-4, in addition to the above analysis, there are some significant differences in the use of criteria. Cancer SIGVAR assigns PM5 to 114 LP/P missense variants, while InterVar did not. Among them, 38 variants are assigned PM5_Strong, and the relative consistency of classification results between Cancer SIGVAR and ClinVar is 100%. InterVar interpreted 8 of the 38 variants as VUS, and the relative consistency with ClinVar was only 78%. For the PM1 criteria, the setting of each parameter is stricter in Cancer SIGVAR, and the functional domain contained only pathogenic or likely pathogenic variants without benign or likely benign variants in these genes.

For the B/LB category of ClinVar, the consistency in the automated interpretation of Cancer SIGVAR is lower than that of InterVar (Figure 3A). As shown in Figure 3D, the differences in the criteria used by the two tools are mainly BS1, BS2, BP1, BP4, and BP6. For the VUS category of ClinVar, 16.87% of the variants were interpreted as LB/B by InterVar, and the main reason is the use of BS2, BP1, and BP4 (Figure 3A-B). Considering the characteristics of cancer-susceptibility genes, Cancer SIGVAR is more rigorous in the use of BS1, BS2 and BP1. BS2 is only applied to *NF1*/*NF2*/*TSC1*/*TSC2* in 97 cancer-susceptibility genes, which is autosomal dominant, and the penetrance rate is extremely high. Among 26,065 ClinVar variants, we screened for a total of 1284 variants (for types, see Supporting Information Figure S3) within the applicable range of BS2. The statistical results show that Cancer SIGVAR assigns 499 variants (38.86%) as BS2, while InterVar assigns only 43 of them (3.35%) and nine other variants (one heterozygote or undocumented). In automated interpretation, Cancer SIGVAR’s assignment conditions based on gnomAD are stricter, and gnomAD has a larger and more comprehensive population base than the 1000 Genome Database used by InterVar; thus, it has a higher and more accurate BS2 assignment rate within the applicable range. This further illustrates the importance of considering the comprehensiveness of the database in automated interpretation. To further analyze the impact of different automated scoring of BS2 on the interpretation of variants, and as far as possible to eliminate the interference of other criteria, we screened 429 B/LB variants (for types, see Supporting Information Figure S4) from the above 1284 variants. The results showed that Cancer SIGVAR assigned BS2 to 246 of these variants (57.34%), while InterVar assigned BS2 to the same 24 variants (5.59%) and the other 5 variants (only one heterozygote in the gnomAD database). For the five InterVar-specific variants, the interpretation results of Cancer SIGVAR and InterVar are completely consistent with those of ClinVar. For the 222 variants assigned as BS2 only by Cancer SIGVAR, the interpretation consistency with ClinVar is shown in Figure 5A. The relative consistency rate of Cancer SIGVAR is 94.59% (210/222), while that of InterVar is 88.29% (196/222). The results show that Cancer SIGVAR’s automated assignment of BS2 is more accurate, which further improves the accuracy of the automated interpretation of variants. Moreover, compared to InterVar, Cancer SIGVAR was able to reduce the proportion of VUS to approximately 53.85% (from 26 to 12 VUS, Figure 5B).

**Figure 5.**
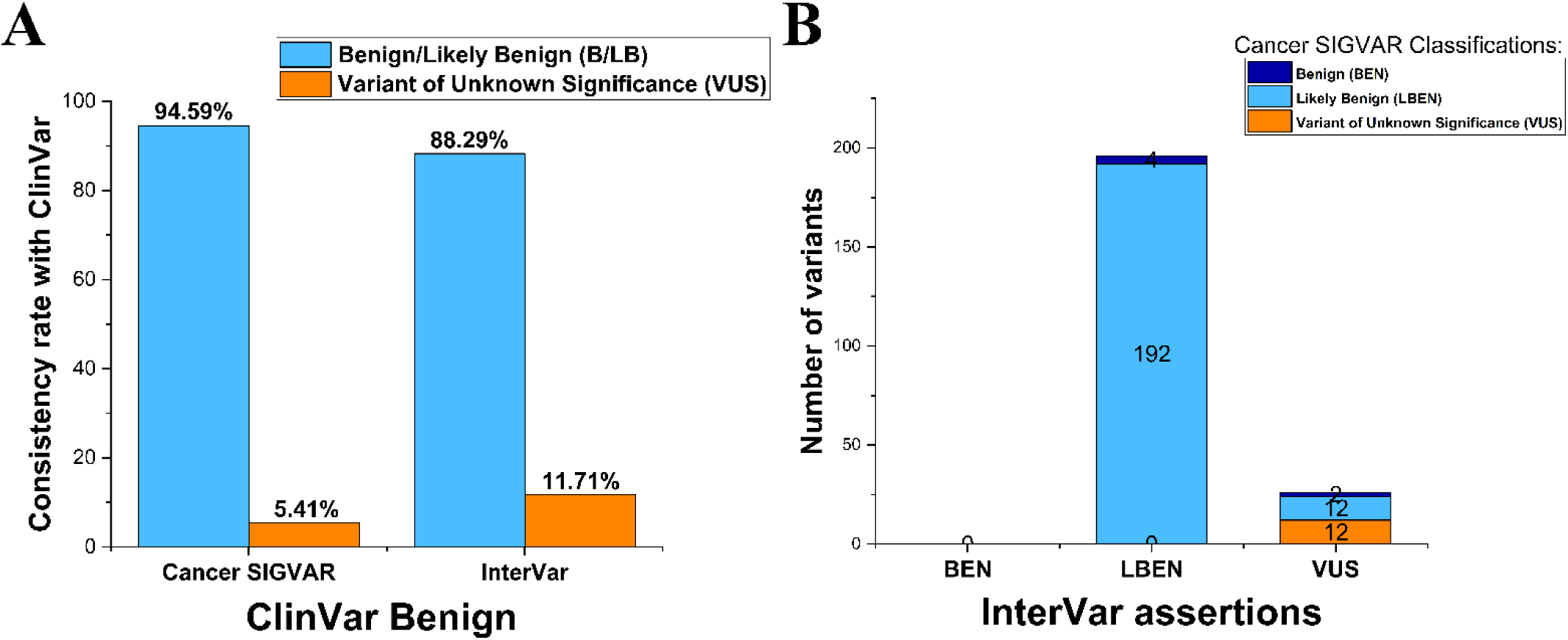
(A). Performance comparison between Cancer SIGVAR and InterVar on 222 ClinVar benign variants. (B). Comparison of InterVar and Cancer SIGVAR classifications for 222 variants.

For PP2 and BP1, a survey found 87 genes out of 97 cancer-susceptibility genes have missense mutation records in the ClinVar database, and 4 of them (*HOXB13*, *CDK4*, *CDK12*, *RAD54L)* satisfy > 80% in pathogenic variants and are missense variants. However, the number of pathogenic variants in the filtered ClinVar database is ≤1, and the amount of data is insufficient. Therefore, the assignment of PP2 criteria should be done with caution if not obtaining more data support. Similar to PP2, considering the reliability and total number of pathogenic variants in the ClinVar database, we are more careful when assigning BP1. As detailed in Supporting Information Table S1, the thresholds are stricter and set according to different modes of inheritance, so the use of BS1 requires more caution. These results further illustrate that Cancer SIGVAR is more accurate in the assignment of criteria and affects the interpretation of variants after the optimization of interpretation rules.

### Logic optimization in Cancer SIGVAR

The consistency in interpretation results is not only due to the adaptation and optimization of the above rules for criteria but also a small part due to the optimization of conflict handling rules. ACGS points out that when PVS1 is used for the interpretation of canonical splice variants, PP3 cannot be used at the same time; otherwise, it will cause the dual use of evidence (Karczewski et al., 2019). We have extended this rule to apply to all types of variants and set logic optimization processing; that is, PP3 should not be used when PVS1 is present. Take a nonsense variant as an example: *MSH3* NM_002439.4 c.1360C>T p. Arg454* (chr5-80021291-CT). According to the results of InterVar, the variant satisfies LOF pathogenicity, and GERP ++ _ RS predicts it as a deleterious variant. Therefore, InterVar assigns PVS1 and PP3. However, Cancer SIGVAR sets up the optimization logic processing for a coexisting conflict between different evidence of the same type, and only the PVS1 criteria are given for this variant. In the LP/P category of ClinVar (Figure 3), InterVar automatically interprets many LP variants as P. Further analysis finds that the difference in interpretation is not only due to the difference in the assignment conditions of PP3 but also due to the simultaneous use of PVS1 and PP3 criteria. InterVar had 1,573 variants applying both PVS1 and PP3 criteria, and all were interpreted to be P. Among the LP category of ClinVar above, 303 variants were interpreted as P by InterVar, and 274 of these used the PVS1 and PP3 criteria simultaneously. The logic optimization between the same type of criteria avoids the overlapping use of evidence and improves the interpretation consistency of Cancer SIGVAR.

When pathogenic and benign criteria coexist, InterVar takes measures to accumulate these pathogenic and benign criteria and then makes a two-way tradeoff. If the two types of evidence are simply superimposed, the extreme contradiction and the difference in the strength of the evidence may be ignored, which will cause the interpretation to lack objectivity. Therefore, Cancer SIGVAR further perfects the classification rules and sets the tolerance value of the criteria. For example, when the criteria combination of PS3, BS3, and BP5 appears, InterVar will classify the variant as likely benign, and Cancer SIGVAR will interpret it as VUS. It is necessary to consider that the possibility of this variant having a significant impact on protein function cannot be ruled out. If such a combination is directly classified as likely benign, there is a risk of false negatives. In another example, when the evidence combination of PS1, PM1, PM2, BP1, BP4 appears, InterVar interprets it as VUS, and Cancer SIGVAR interprets it as likely pathogenic. For variants that already have the same codon pathogenesis record (PS1), the probability of software predicting harmless (BP4) is low. Considering that the former criteria are more reliable than the latter, it tends to take PS1 criteria. Comprehensive analysis of those criteria combinations should be classified as likely pathogenic.

### Interpretation accuracy of Cancer SIGVAR semiautomatic based on all 48 criteria

To evaluate the variant interpretation accuracy of the entire process of Cancer SIGVAR, we selected two artificial interpretation cancer variant datasets (Datasets 3 and 4) that are not public on the website (see Materials and Methods for details). Dataset 3 has 731 cancer variants from the National Institutes for Food and Drug Control. In addition to the results from the automated interpretation, a third step, manual supplementary analysis of the entire interpretation process, is added. Consistency with the manual interpretation results is shown in Table 4. The consistency of Cancer SIGVAR’s automated interpretation of P/LP categories was 97.15%, and the consistency of interpretation was increased to 99.29% after adding the third step of manual supplementary criteria. The consistency of B/LB variant interpretation increased from 81.93% of automated interpretation to 100% of the entire process. The interpretation consistency of all variants in the entire process is 98.91%. The interpretation consistency of different types of variants is shown in Figure 6. To better test the accuracy of Cancer SIGVAR’s interpretation of variants with more semiautomatic criteria and conflicting evidence, we selected 180 variants in the in-house database of BGI to include as many criteria as possible. The results show that the consensus rate in the automated interpretation of Cancer SIGVAR is 63.33% (114/180). After adding the semiautomatic criteria and the manual supplementary criteria in the third step, the interpretation rate of Cancer SIGVAR was 97.78% (176/180). The frequency of the use of criteria is shown in Figure 7. The interpretation of a few variants in the entire process is inconsistent, and 100% consistency is not reached because of differences in database updates and in the frequency database. The results show that the entire process followed by Cancer SIGVAR has a high accuracy in variant interpretation.

**Table 4.** Comparison of Variant Interpretation consistent with China Institutes database by Automatic and Manual Adjusted of Cancer SIGVAR.

| Clinical Significance | China Institutes database | Cancer SIGVAR (Automated Interpretation) | Cancer SIGVAR ( Interpretation after Manual Adjustment) |
| --- | --- | --- | --- |
| Pathogenic or likely pathogenic | 421 | 409 (97.15%) | 418 (99.29%) |
| Benign or likely benign | 83 | 68 (81.93%) | 83 (100.00%) |
| VUS | 227 | 222 (97.80%) | 222 (97.80%) |
| Sum of five tiers | 731 | 699 (95.62%) | 723 (98.91%) |

**Figure 6.**
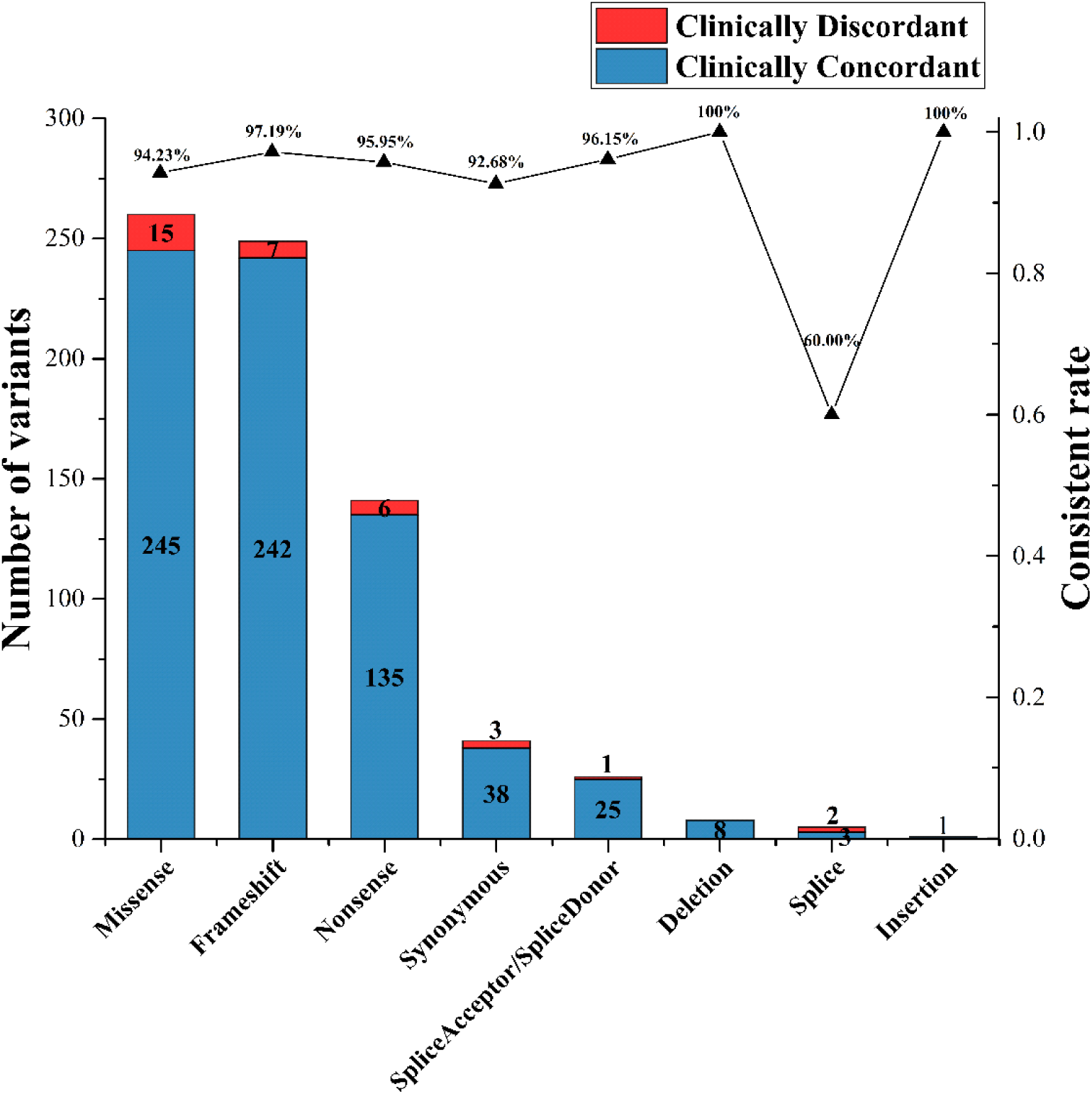
Consistency of interpretation of different types of variant, which was interpreted by the entire process of Cancer SIGVAR

**Figure 7.**
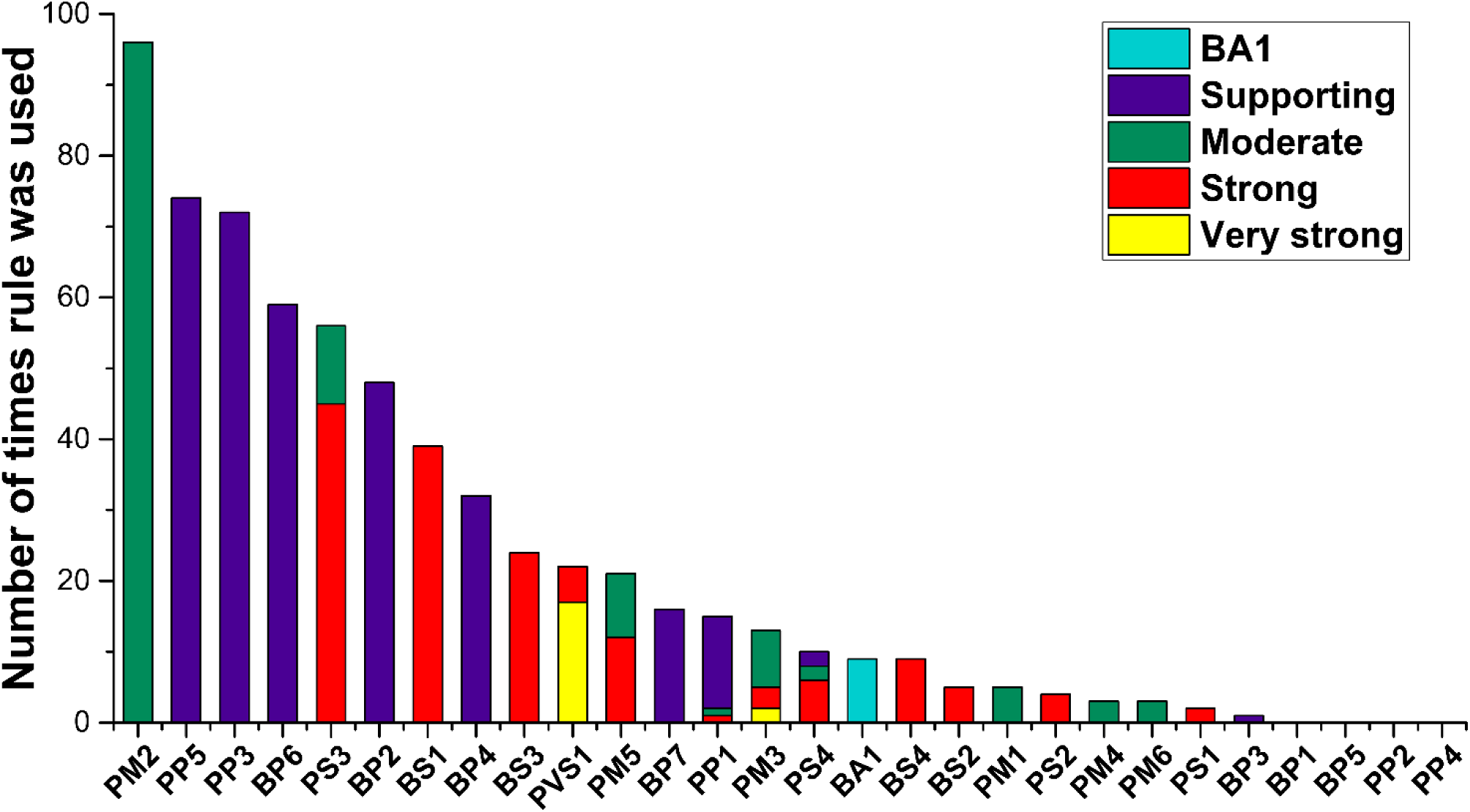
Summary of criteria that were applied to the 180 validation variants

## Discussion

The ACMG-AMP guidelines set the standard for the interpretation of genetic variants in clinical practice. As the ClinGen SVI Working Group continues to refine the use of criteria and optimize disease specificity, interpretation consistency has been improved, but it has also increased the complexity and error-proneness of manual interpretation. Therefore, it is necessary to develop an automated interpretation tool for hereditary cancer combined with the latest guidelines. For this purpose, we have developed Cancer SIGVAR, a semi-automated hereditary cancer interpretation tool.

We integrated different omics data sources, such as RefSeq, InterPro, UniProt, OMIM, Genereview, Gene Condition, OrphaNet, ClinVar, and customized cancer gene-disease related knowledge bases, including gene-phenotype relationships, gene functional domains, gene variation pathogenic mechanism (LOF), protein repeat region, NMD prediction library, etc. We implemented different strategies to trigger the criteria and built a complete library of pathogenic/benign variants. The integration of these resources also emphasizes the importance of sharing data through a centralized database. Second, we adopted the latest adaptation of the guidelines proposed by the ClinGen Working Group (Gelb et al., 2018; Mester et al., 2018; Oza et al., 2018), expanding 28 criteria to 48 criteria and refining its rules of use. Third, we set up logical processing according to the latest guide adaptation, reducing the reuse of similar evidence, and setting the tolerance value to optimize the classification rules. In addition, we proposed some extensions or refinement of the criteria, such as BP2, PM2 and PM4.

We determined the extremely low frequency threshold of PM2 by statistical analysis of the pathogenic variants in the database. The detailed statistical analysis process is shown in Supporting Information Figure S5. In the future, thresholds can be determined based on more databases for analysis so that criteria can be as accurately assigned as possible. We propose a refinement of the BP2 criteria. The results for the proportion of homozygotes detected in different pathogenic variants in the CLINVITAE and ClinVar databases are shown in Supporting Information Figure S6. For the dominant gene in the hereditary cancer panel, the pathogenic probability of the homozygous variant is very low, which supports the assignment of BP2 criteria. In addition, a similar refinement of BP2 criteria has also been specified to individual genes of interest by ClinGen Expert Panel Groups, including *CDH1* (Lee et al., 2018) and *RUNX1* (Luo et al., 2019). For some unidentified variants, due to a lack of necessary supporting data, the refinement of BP2 criteria can make the clinical significance clearer and to some extent reduce the proportion of unidentified variants. According to the PVS1 ACMG/AMP criterion by ClinGen (Abou Tayoun et al., 2018), if variants of the LOF gene, such as frameshift or nonsense mutations, cause a protein length loss of more than 10%, PVS1_Strong should be applied. Considering that the in-frame deletion variant may also cause the protein loss length to exceed 10%, to maintain the consistency of the criteria strength, we have extended the PM4 criteria. For the LOF gene, PM4_Strong will be assigned if the protein length loss of in-frame deletion exceeds 10%. This refinement may make the interpretation of some variants more accurate.

Cancer SIGVAR comprehensively considers the use of criteria for different types of variants. Fully automated interpretations based on 21 criteria show an average interpretation consistency of 98.95% on two different benchmark datasets (ClinVar and CLINVITAE). Compared with the similar tool InterVar, the results show that the interpretation of Cancer SIGVAR is more consistent, especially for pathogenic variants. For PVS1 criteria, Cancer SIGVAR has a more accurate evidence level assignment for fully considering the genetic structure and pathophysiological mechanism (such as NMD or alternative splicing), which greatly improves the consistency of the pathogenic interpretation results. In the case of NMD, in particular, the consistency of pathogenic interpretation was improved by 54.42% compared to InterVar. In addition, we analyzed significant differences in criteria assignment. The differences in criteria PS1, PP5, and BP6 depend on matching with established pathogenic variants, which confirms the importance of laboratory management and maintenance of a truly pathogenic and benign variant pool. The difference in criteria BS1 depends on the threshold of the population database. It cannot be so high that it is never reached, nor too low to be overused, as either situation will lead to the misclassification of variants (Oza et al., 2018). The threshold of BS1 in Cancer SIGVAR is set separately for recessive and dominant, which is stricter and avoids overuse. The difference in criteria BS2 confirms the importance of considering the scope of application of the criteria. These analyses highlight the flexibility of the ACMG-AMP criteria implementation (Hoskinson, Dubuc, & Mason-Suares, 2017).

We want to emphasize that the Cancer SIGVAR interpretation includes 2 processes. The first is to output preliminary automated interpretation results based on 21 automated criteria. The second is based on 21 automated criteria and adds 27 criteria through semiautomatic or manual interpretation based on considerations of disease mechanism, co-segregation, disease and gene specificity, comprehensively adjusting the output of similar criteria and providing a final interpretation. In general, automated tools can reduce the complications and error-proneness of searching. Second, semiautomated scoring improves the efficiency and accuracy of the criteria level calculation. Finally, experts can check the output of automated criteria and additional inputs to obtain the most accurate interpretation of genetic variants. We verified the entire process through a total of 911 variants of two in-house benchmark datasets. The average classification consistency of Cancer SIGVAR was 98.35%.

Cancer SIGVAR also has limitations. First, this tool complies with the requirements for personalized interpretation of cancer genes but currently only applies to 97 genes of hereditary cancer. Future versions will expand to include more genes of interest. Considering this, we have designed a unified configuration mode with high configurability. Later, stages can be expanded according to similar methods and can be customized according to the criteria constraints and classification rules of different diseases. Second, Cancer SIGVAR is currently limited to the interpretation of SNVs and insertions/deletions (INDELs). CNV interpretation rule investigations have been carried out, and future versions will try to integrate the ability to analyze CNV and other genetic variants. Finally, the ClinGen Working Group is revising the criteria use and classification rules for the ACMG/AMP guidelines, and we will continue to develop Cancer SIGVAR so that it can be used quickly when these guidelines are updated.

In summary, we developed an automated cancer interpretation tool, Cancer SIGVAR, based on the 2015 ACMG-AMP guidelines and various optimized adaptations by ClinGen Sequence Variation Interpretation Expert Groups. The criteria were expanded from 28 to 48. Cancer SIGVAR can achieve an automated scoring of 21 criteria (43.8%), automatically generate a preliminary interpretation, then achieve a semiautomatic scoring of 17 criteria (35.4%) and allow manual adjustment of additional criteria to arrive at a definitive final interpretation. Cancer SIGVAR improves the accuracy of the automated interpretation of hereditary cancer, promotes the clinical evaluation of variants, and lays the foundation for fully automated interpretation and accurate diagnosis and treatment of cancer.

## Supplementary Material

Refer to Web version on PubMed Central for supplementary material

## Acknowledgments

Hong Li, Shuixia Liu and Quanlei Zeng conceived the study. Yulan Chen, Yun Xiong, Shuixia Liu and Yi Zhang implemented the software. Shuangying Wang wrote the first draft of the manuscript. Hong Li, Shuixia Liu, Quanlei Zeng, Ting Fang, Yu Zhang, Yun Xiong, Yulan Chen and Yi Zhang provided critical edits and feedback. The following authors have contributed to the interpretation rules and implement of Cancer SIGVAR: Ying Zhou, Yi Zhang, Kaiyue Wang, Zhangwei Yan, Cuicui Qiang, Meng Xu, Xianghua Chai. The work was overseen by Yun Xiong, Yuying Yuan and Hongyun Zhang. Ming Huang provides help for data testing performance. We would like to acknowledge National Institutes for Food and Drug Control staff, in particular Jie Huang, for the support of data testing performance

