## Supporting Information for "Cancer SIGVAR: A semi-automated interpretation tool for germline variants of hereditary cancer-related genes"

#### SUPPLEMENTAL FIGURES AND METHODS

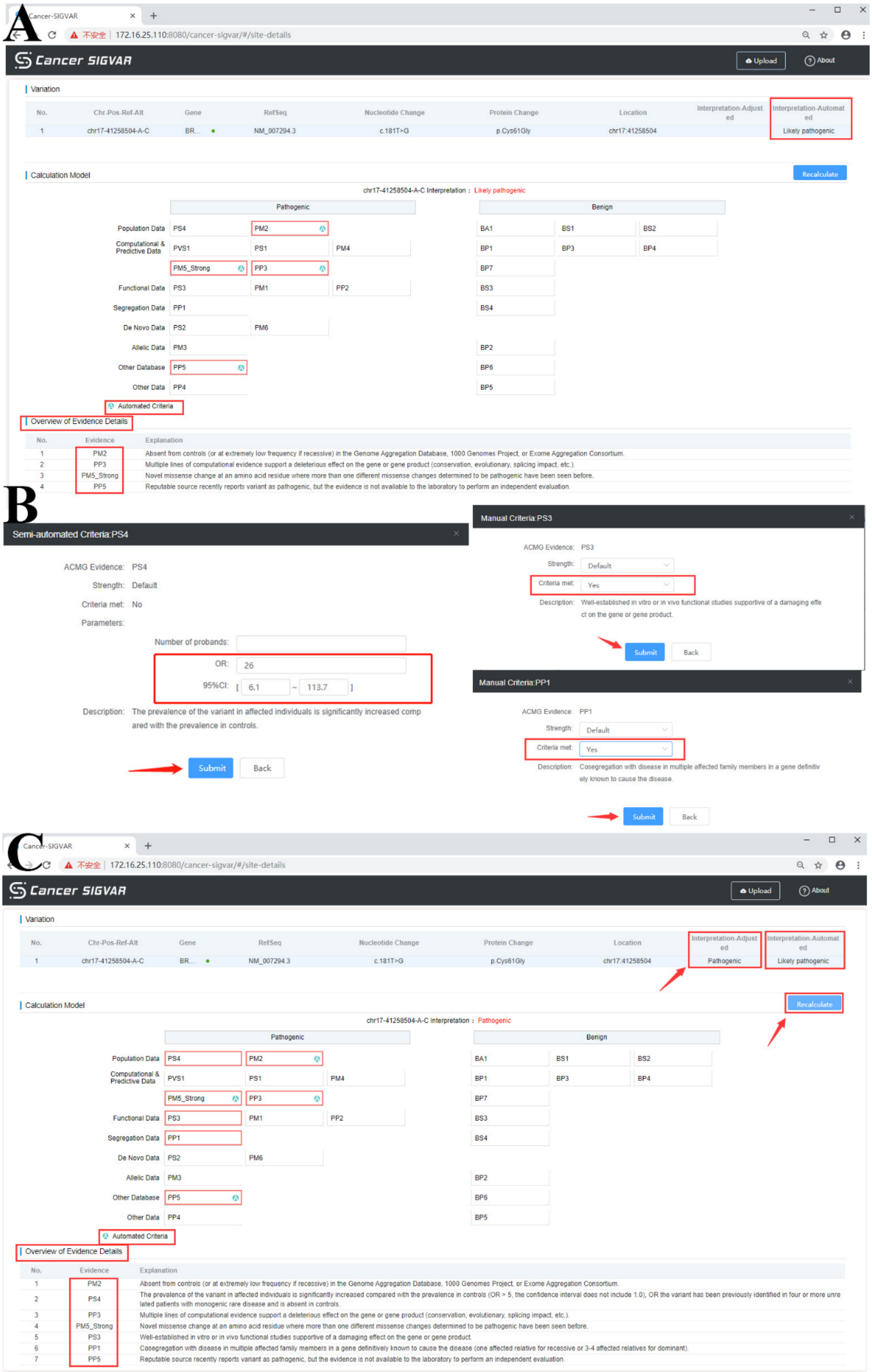


**Figure S1. Web interface display of Cancer SIGVAR interpretation process. (A) Automatic interpretation of genetic variant. (B) manual adjustment. manually entering parameters for semi-automated scoring and manual assignment. (C) Click “Recalculate”, then the reinterpret results and full list of criteria are shown.**

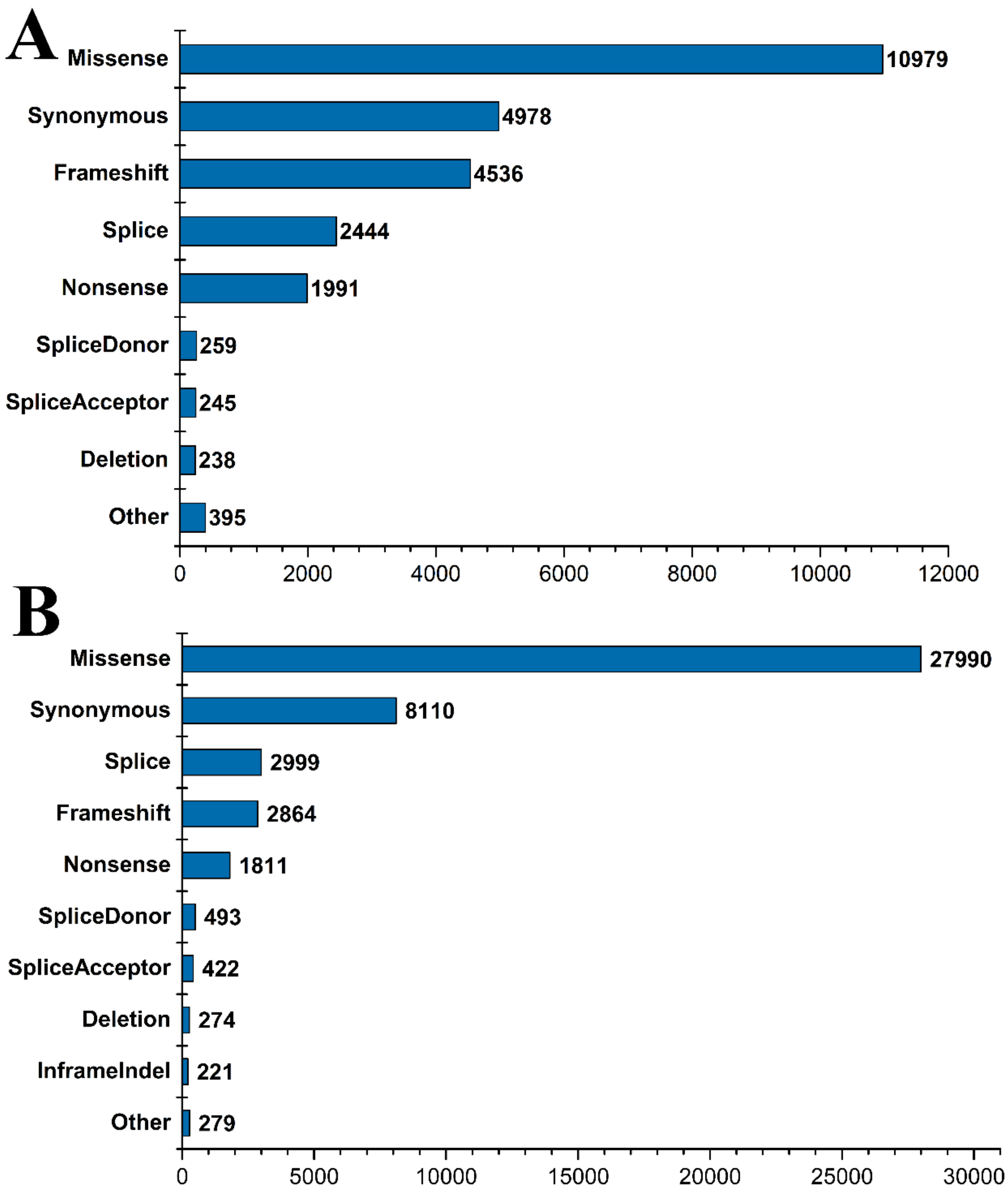


**Figure S2. (A). Types of 26065 variants. selected from the ClinVar dataset. (B). Types of 45463 variants selected from the CLINVITAE dataset.**

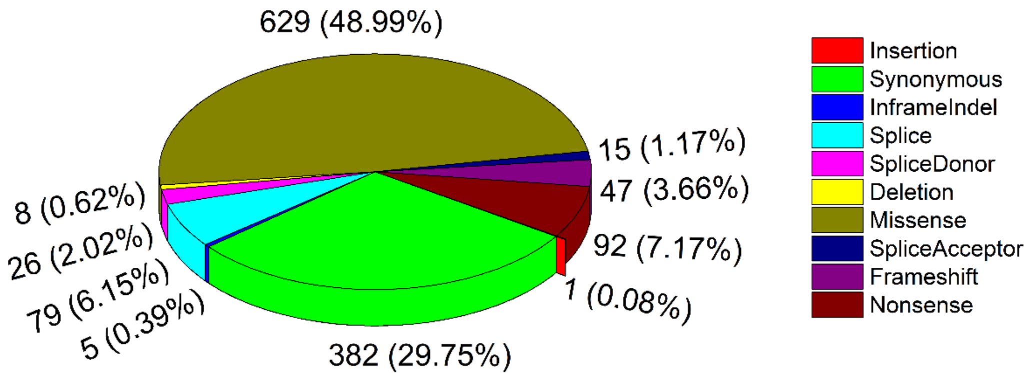


**Figure S3. Types of alterations for 1284 variants selected for validation.**

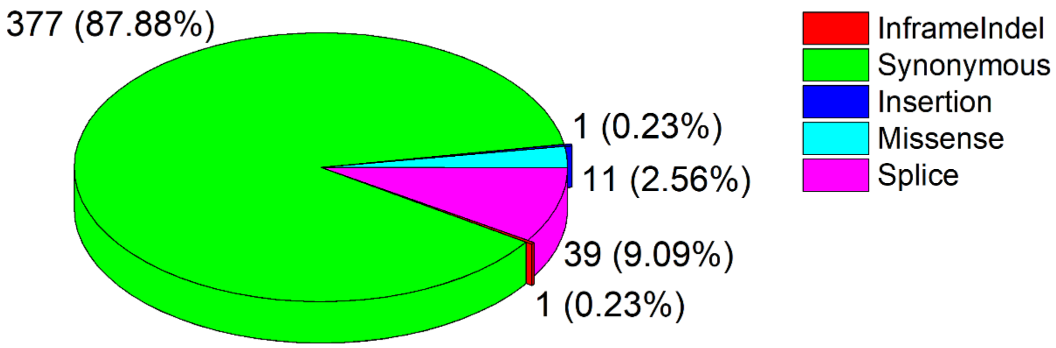


**Figure S4. Types of alterations for 429 benign variants selected for validation.**

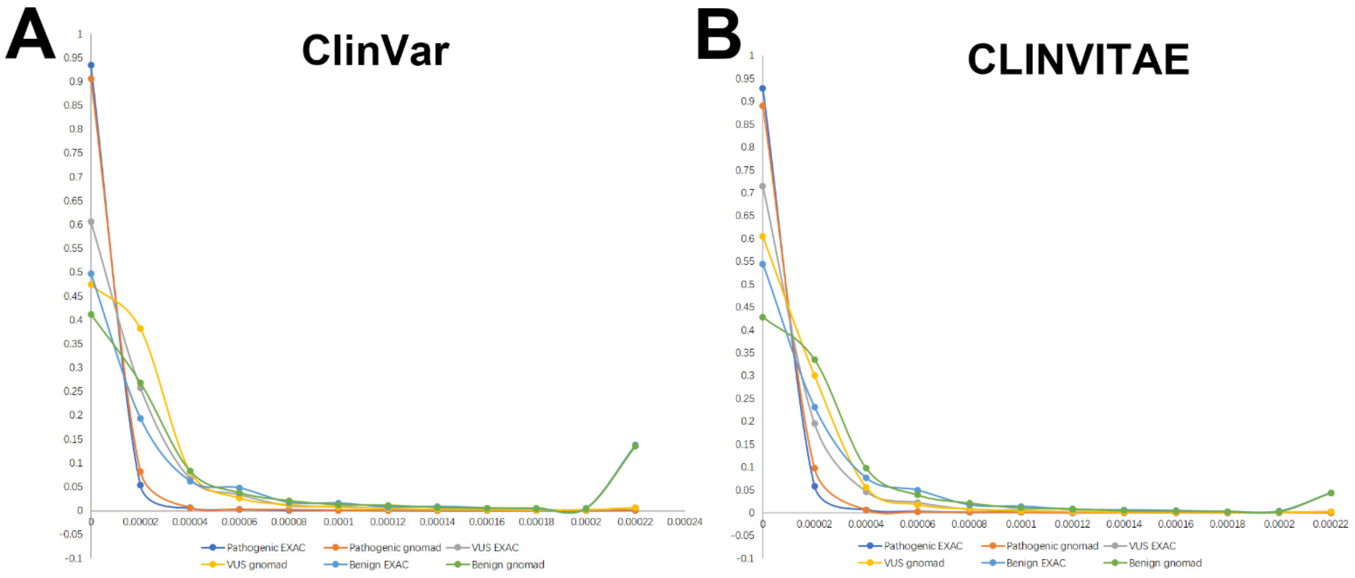


**Figure S5. Frequency distribution of various pathogenic variants in ClinVar (A) and CLINVITAE (B)**

We selected 32,661 variants with the review status ≥ 2 stars of genetic tumors gene panel in ClinVar database, which involved a total of 10,154 pathogenic variants, 12,610 unidentified variants, and 9897 benign variants. Count the number of variants of different clinical significance in gnomAD and ExAC in different frequency ranges and calculate the proportion in different frequency ranges. The results are shown in Figure S5A. The proportions of pathogenic variants that satisfy MAF ≤ 0.00008 in gnomAD and ExAC are 99.79% and 99.68%, respectively. Analysis of 45,399 variants (5711 pathogenic variants, 30,284 unidentified variants, and 9404 benign variants) in the hereditary tumor genes in the CLINVITAE database. The variation distribution of different frequency ranges is shown in Figure S5B. The pathogenic variation with MAF ≤ 0.00008 was 99.84% and 99.69% in the gnomAD and ExAC, respectively. If a variant is absent from (or below the expected carrier frequency if recessive) a large general population or a control cohort (>1,000 individuals), it can be considered a moderate piece of evidence for pathogenicity (PM2) (Richards et al., 2015). We set the threshold of PM2 as 0.00008. On the one hand, it meets the requirements below the expected frequency, and on the other hand, the average probability of pathogenic variation is 99.75%, which meets the threshold condition. Rare variants with frequencies below 0.00008 can be used as evidence to support pathogenicity.

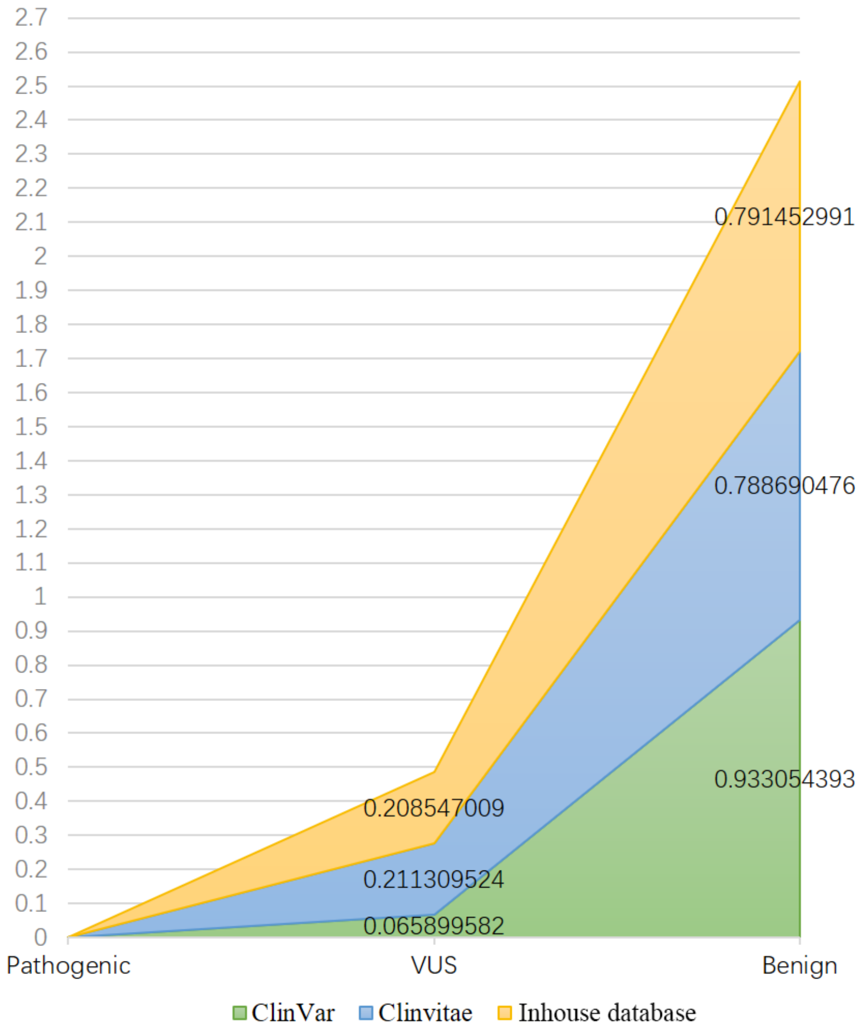


**Figure S6. Various clinically significant distributions of homozygotes in ClinVar, CLINVITAE, and inhouse databases.**

Statistics the various pathogenic variant distributions of homozygotes in ClinVar, CLINVITAE and inhouse database are shown in Figure S6. The proportions of different clinical significance of homozygous variant in the three databases are basically the same. The proportion of homozygous present in the pathogenic variants is close to 0. The percentage of homozygotes detected increases as variants tend to benign. For dominant genes, homozygotes are detected in the gnomAD population frequency library, and the probability that the homozygotes will eventually be judged to be benign is close to 80%, which satisfies supporting benign evidence. Therefore, we propose the expansion of BP2. For hereditary tumors detect panel-related dominant gene variants, if homozygotes are detected, the variant supports benign evidence and assigns BP2.

### SUPPLEMENTARY TABLES

#### Table S1. ACMG-AMP criteria and their implementation in Cancer SIGVAR.

| category | Criteria | Original ACMG-AMP description | Specification | Reference | Cancer SIGVAR Implementation |
|---|---|---|---|---|---|
| Computational & Predictive Data | PVS1 | Null variant in genes with LOF (loss of function) as known mechanism of disease | Automated Scoring strength PVS1, PVS1_Strong, PVS1_Moderate | (Abou Tayoun et al., 2018) | We consider the determination of LOF genes and variant type-specific considerations, gene structure and pathophysiologic mechanisms, and refer to the strength level rules of PVS1 proposed by ClinGen SVI. (See PVS1 in Figure 1 for details). |
| | PS1 | Same amino acid change as a previously established pathogenic variant | Automated Scoring NA | (Mester et al., 2018; Oza et al., 2018) | Same amino acid change as a previously established pathogenic variant OR same nucleotide position as a previously established pathogenic variant that affects splicing |
| | PM4 | Protein length changes as a result of in-frame deletions/insertions in a nonrepeat region or stop-loss variants | Automated Scoring strength PM4, PM4_Strong | (Abou Tayoun et al., 2018; Karczewski et al., 2019) | PM4_Strong: length of amino acid deletion > 10% length of wildtype protein; PM4: length of amino acid deletion ≤10% length of wildtype protein |
| | PM5 | Novel missense change at an amino acid residue where a different missense change determined to be pathogenic have been seen before | Automated Scoring strength PM5, PM5_Strong | (Gelb et al., 2018) | PM5_Strong: ≥ 2 different pathogenic missenses; PM5: 1 different pathogenic missense |
| | BP1 | Missense variant in a gene for which primarily truncating variants are known to cause disease | Automated Scoring NA | (Li & Wang, 2017) | Missense variant in a gene which truncating variants (> 80% in pathogenic variants) are known to cause disease. BP1 was caution to apply |
| | BP3 | In-frame deletions/insertions in a repetitive region without a known function | Automated Scoring NA | - | Variant is in-frame deletion/insertion, but overlaps a repetitive region |
| | PP3 | Multiple lines of computational evidence support a deleterious effect on the gene or gene product (conservation, evolutionary, splicing impact, etc.) | Automated Scoring NA | (Karczewski et al., 2019) | Variant is Missense and "pathogenic" for SIFT, PolyPhen-2, Mutation Taster and dbscSNV; Variant is Splice and "pathogenic" for dbscSNV; Variant is synonymous and "pathogenic" for dbscSNV, GERP and DANN. PP3 cannot be applied in combination with PVS1. |
| | BP4 | Multiple lines of computational evidence suggest no impact on gene or gene product | Automated Scoring NA | | Variant is Missense and "benign" for SIFT, PolyPhen-2, Mutation Taster and dbscSNV; Variant is Splice and "benign" for dbscSNV; Variant is synonymous and "benign" for dbscSNV, GERP and DANN. BP4 cannot be applied in combination with PVS1 |
| | BP7 | A synonymous variant for which splicing prediction algorithms predict no impact to the splice consensus sequence nor the creation of a new splice site and the nucleotide is not highly conserved | Automated Scoring NA | - | Variant is synonymous and "benign" for dbscSNV |

| | | | | | |
|---|---|---|---|---|---|
| Population_data | PS4 | The prevalence of the variant in affected individuals is significantly increased compared with the prevalence in control subjects | Semi-automatic Scoring strength PS4, PS4_Moderate, PS4_Supporting | (Luo et al., 2019; Mester et al., 2018) | The Case accumulation is only applicable to the NF1 / NF2 / TSC1 / TSC2 gene variants, assign PS4 and its strength level based on the rules proposed by ClinGen's RASopathy Expert. For all variants, we use the odds ratio (OR) to assign PS4 and set downgrading to PS4_Supporting when 3≤OR≤5 |
| | PM2 | Absent from controls (or at extremely low frequency if recessive) | Automated Scoring NA | (Chen et al., 2016; Siegel, Miller, & Jemal, 2017) | Use five population databases. both AF of gnomAD and ExAC ≤ 0.00008 and all AF <0.005 (AD in OMIM) or all AF <0.01 (AR in OMIM) |
| | BA1 | Allele frequency is >5% in population databases such as ESP, 1000G, or ExAC | Automated Scoring NA | - | Any allele frequency is >5% in five population databases |
| | BS1 | Allele frequency is greater than expected for disorder | Automated Scoring NA | (Chen et al., 2016; Siegel, Miller, & Jemal, 2017) | The threshold for BS1 based on the total incidence of cancer. use five population databases. all AF ≤ 0.05 and ≥ 0.005 (AD in OMIM) or all AF ≤ 0.05 and ≥ 0.01 (AR in OMIM) |
| | BS2 | Observed in a healthy adult individual for a recessive (homozygous), dominant (heterozygous), or X-linked (hemizygous) disorder, with full penetrance expected at an early age | Automated Scoring NA | - | Full penetrance expected at an early age (NF1 / NF2 / TSC1 / TSC2 in 97 cancer-susceptibility genes) and ≥ 2 homozygotes/heterozygotes/hemizygotes are observed in the healthy adult come from gnomAD |
| Segregation Data | PP1 | Cosegregation with disease in multiple affected family members in a gene definitively known to cause the disease | Manual Scoring Strength PP1, PP1_Moderate, PP1_Strong | (Lee et al., 2018; Oza et al., 2018) | If the genetic variant co-segregated with disease (AD), it was assigned according to the rule in previous study by the number of meiotic divisions (excluding probands) and family. When the inheritance is AR, it was assigned according to the rule in previous study by the LOD score |
| | BS4 | Lack of segregation in affected members of a family | Manual Scoring NA | - | Lack of co-segregation in probands |
| Denovo Data | PS2 | De novo (maternity and paternity confirmed) in a patient with the disease and no family history | Semi-automatic Scoring strength PS2, PS2_Very strong, PS2_Moderate, PS2_Supporting | (Oza et al., 2018) | Point value is according to the occurrence(s) and phenotypic consistency of de novo. total points ≥ 4 (PS2_Very_strong); 2≤total points<4 (PS2); 1≤total points<2 (PS2_Moderate); total points =0.5 (PS2_Supporting) |
| | PM6 | Assumed de novo (without confirmation of paternity and maternity) in a patient with the disease and no family history | Semi-automatic Scoring strength PM6, PM6_Very strong, PM6_Strong, PM6_Supporting, PM6_Supporting_0. | | Point value is according to the occurrence(s) and phenotypic consistency of de novo. total points ≥ 4 (PM6_Very_strong); 2≤total points<4 (PM6_Strong); 1≤total points<2 (PM6); total points =0.5 (PM6_Supporting) |

| | | | | | |
|---|---|---|---|---|---|
| | | | 5 | | |
| Functional Data | PS3 | Well-established in vitro or in vivo functional studies supportive of a damaging effect on the gene or gene product | Manual Scoring strength PS3, PS3_Moderate | (Oza et al., 2018) | ClinGen HL-EP have pointed out that when the reliability of functional experiments is low, the intensity of PS3 can be reduced to a medium or support level. Based on the results of functional experiments in the literature, we assign PS3_Moderate if the function experiment weak support |
| | PM1 | Located in a mutational hot spot and/or critical and well-established functional domain without benign variation | Automated Scoring NA | - | Variant within a known hotspot domain and well-established functional domain without benign variation |
| | PP2 | Missense variant in a gene that has a low rate of benign missense variation and in which missense variants are a common mechanism of disease | Manual Scoring NA | (Li & Wang, 2017) | According to intervar's implementation rules, missense variant in a gene that has a low rate of benign missense variant (<10%) and in which missense variants (> 80%) are pathogenic variants |
| | BS3 | Well-established in vitro or in vivo functional studies show no damaging effect on protein function or splicing | Manual Scoring NA | - | Well-established functional studies with no damaging effect |
| Allelic Data | PM3 | For recessive disorders, detected in trans with a pathogenic variant | Semi-automatic Scoring strength PM3_Very strong, PM3_Strong, PM3, PM3_Supporting, PM3_Supporting_0.5 | (Oza et al., 2018; Karczewski et al., 2019) | We refer to the PM3 rules of ClinGen HL-EP, and established a scoring system. Points per proband is seen in Supporting Information TableS2. The final level of criteria was determined by the total points. |
| | BP2 | Observed in trans with a pathogenic variant for a fully penetrant dominant gene/disorder or observed in cis with a pathogenic variant in any inheritance pattern | Automated Scoring NA | (Luo et al., 2019; Nykamp et al., 2017) | Homozygotes are detected in dominant gene variants, detailed statistical analysis process seen in Supporting Information Figure S3 |
| Other Data | PP5 | Reputable source recently reports variant as pathogenic, but the evidence is not available to the laboratory to perform an independent evaluation | Automated Scoring NA | (Biesecker & Harrison, 2018) | To avoid abuse, we only apply PP5 when the conclusion in ClinVar as pathogenic, not likely pathogenic. Using BP6 is also more rigorous. In addition, only one PP5 or BP6 will be output at most |
| | BP6 | Reputable source recently reports variant as benign, but the evidence is not available to the laboratory to perform an independent evaluation | Automated Scoring NA | | |
| | PP4 | Patients phenotype or family history is highly specific for a disease with a single genetic etiology | Manual Scoring NA | - | - |

| | BP5 | Variant found in a case with an alternate molecular basis for disease | Manual Scoring NA | - | - |
|---|---|---|---|---|---|

**Table S2. Points per proband for scoring variants that are observed in trans (PM3 rules)**

| Classification/zygosity of other variant | Known in trans |
|---|---|
| Pathogenic/Likely pathogenic | 1.0 |
| Homozygous occurrence<br>(Max points = 1.0) | 0.5 |
| Rare uncertain significance variant on other allele,<br>OR Homozygous occurrence due to consanguinity,<br>(Max point = 0.5) | 0.25 |

**Table S3 Rules and logic of automated pathogenicity classification**

| Number | Interpretation | PVS | PS | PM | PP | BS | BP | BA |
|---|---|---|---|---|---|---|---|---|
| R1 | Benign | n | n | n | n | n | n | ≥1 |
| R2 | Benign | 0 | 0 | 1 | [0~1] | ≥2 | n | 0 |
| R3 | Benign | 0 | 0 | 0 | n | ≥2 | n | 0 |
| R4 | Likely Benign | 0 | 0 | 2 | [0~1] | ≥2 | n | 0 |
| R5 | Likely Benign | 0 | 0 | 1 | [2~3] | ≥2 | n | 0 |
| R6 | Likely Benign | 0 | 0 | 1 | [0~1] | 1 | ≥1 | 0 |
| R7 | Likely Benign | 0 | 0 | 0 | n | 1 | ≥1 | 0 |
| R8 | Likely Benign | 0 | 0 | 1 | [0~1] | 0 | ≥2 | 0 |
| R9 | Likely Benign | 0 | 0 | 0 | n | 0 | ≥2 | 0 |
| R10 | Likely Pathogenic | 1 | 0 | 1 | 0 | 0 | n | 0 |
| R11 | Likely Pathogenic | 0 | 1 | 1 | [0~3] | 0 | n | 0 |
| R12 | Likely Pathogenic | 0 | 1 | 2 | [0~1] | 0 | n | 0 |
| R13 | Likely Pathogenic | 0 | 1 | 0 | ≥2 | 0 | n | 0 |
| R14 | Likely Pathogenic | 0 | 0 | ≥3 | n | 0 | [0~1] | 0 |
| R15 | Likely Pathogenic | 0 | 0 | 2 | ≥2 | 0 | [0~1] | 0 |
| R16 | Likely Pathogenic | 0 | 0 | 1 | ≥4 | 0 | [0~1] | 0 |
| R17 | Pathogenic | ≥2 | n | n | n | 0 | n | 0 |
| R18 | Pathogenic | 1 | ≥1 | n | n | 0 | n | 0 |
| R19 | Pathogenic | 1 | 0 | ≥2 | n | 0 | n | 0 |
| R20 | Pathogenic | 1 | 0 | 1 | ≥1 | 0 | n | 0 |
| R21 | Pathogenic | 1 | 0 | 0 | ≥2 | 0 | n | 0 |
| R22 | Pathogenic | 0 | ≥2 | n | n | 0 | n | 0 |
| R23 | Pathogenic | 0 | 1 | ≥3 | n | 0 | n | 0 |
| R24 | Pathogenic | 0 | 1 | 2 | ≥2 | 0 | n | 0 |
| R25 | Pathogenic | 0 | 1 | 1 | ≥4 | 0 | n | 0 |
| R26 | VUS | Others | | | | | | |

**Table S4. 97 susceptible genes in Cancer SIGVAR.**

| | | | | | | |
|---|---|---|---|---|---|---|
| ALK | APC | ATM | ATR | ATRX | AXIN2 | BAP1 |
| BARD1 | BLM | BMPR1A | BRCA1 | BRCA2 | BRIP1 | CDC73 |
| CDH1 | CDK12 | CDK4 | CDKN1B | CDKN2A | CHEK1 | CHEK2 |
| DMC1 | EME1 | EME2 | EPCAM | EXT1 | EXT2 | FAM175A |
| FANCA | FANCG | FANCI | FANCL | FH | FLCN | GALNT12 |
| GEN1 | HOXB13 | KIT | MAX | MEN1 | MET | MLH1 |
| MLH3 | MRE11A | MSH2 | MSH3 | MSH6 | MUS81 | MUTYH |
| NBN | NF1 | NF2 | NTHL1 | NTRK1 | PALB2 | PDGFRA |
| PHOX2B | PMS1 | PMS2 | POLD1 | POLE | PPP2R2A | PRSS1 |
| PTCH1 | PTCH2 | PTEN | RAD50 | RAD51B | RAD51C | RAD51D |
| RAD52 | RAD54L | RB1 | RBBP8 | RET | SDHA | SDHAF2 |
| SDHB | SDHC | SDHD | SLX1A | SLX4 | SMAD4 | SMARCA4 |
| SPINK1 | STK11 | SUFU | TMEM127 | TP53 | TP53BP1 | TSC1 |
| TSC2 | VHL | WT1 | XPC | XRCC2 | XRCC3 | |

**Table S5. Comparison of Variant Automated Interpretation consistent with CLINVITAE by InterVar and Cancer SIGVAR**

| Clinical Significance | CLINVITAE | InterVar (Automated Interpretation) | Cancer SIGVAR (Automated Interpretation) |
|---|---|---|---|
| Pathogenic or likely pathogenic | 5775 | 4920 (85.19%) | 5316 (92.05%) |
| Benign or likely benign | 9404 | 7855 (83.53%) | 7676 (81.62%) |
| VUS | 30284 | 25277 (83.47%) | 29959 (98.93%) |
| Sum of five tiers | 45463 | 38052 (83.70%) | 42951 (94.47%) |

The results show that the consistency rate of Cancer SIGVAR is 92.05% (5316/5775) for the P/LP category of CLINVITAE, which is higher than that of InterVar, 85.19% (4920/5775). For the B/LB category, the consistency rate of Cancer SIGVAR is 81.62% (7676/9404), which is slightly lower than the 83.53% (7855/9404) of InterVar. The overall consistency of the automated interpretation of Cancer SIGVAR is 94.47% (42951/45463), which is higher than that of InterVar 83.70% (38052/45463).
